# Comparative severity analysis of colitis in C57BL/6 than BALB/c mice: A novel and rapid model of DSS induced colitis

**DOI:** 10.1101/2020.05.07.082669

**Authors:** Sohini Mukhopadhyay, Palok Aich

## Abstract

Colitis is a complex and multifactorial disease with unknown etiology. To understand colitis, a number of models were developed in mouse. Colitis, in these models, was induced using either chemicals or select bacteria. Dextran sulfate sodium (DSS) induced colitis model was used widely, over other models, because of the simpler properties of DSS and similarities of the model with colitis in humans. The available models of DSS induced colitis require either longer time to understand the chronic stages of the disease or a shorter time period to study the acute phase. Currently, no composite models exist that can be used to study the acute, chronic and recovery stages of colitis within a reasonably shorter period of time. In the current study, we have developed a newer model system in two differently immune biased (Th-1 and Th-2) mice strains using DSS. We have developed the DSS model to compare the response in C57BL/6 (Th1 biased) and BALB/c (Th2 biased) mice models. We have standardized the dosage for both C57BL/6 and BALB/c mice with zero mortality rates. Using the model developed we studied the acute, chronic and recovery phases of colitis. The differential responses of C57BL/6 and BALB/c revealed that the disease was more severe in C57Bl/6. BALB/c, on the contrary, recovered from the diseased condition more quickly. The current report will further help to unravel the disease etiology based on immunological background of the host.

## Introduction

The mammalian gastrointestinal tract is continuously exposed to numerous microbial pathogenesis well as food-derived and environmental toxins that could lead to diseases such as Crohn’s disease (CD) and ulcerative colitis (UC) or inflammatory bowel diseases (IBD) (1). IBD patients show flares of remission and relapses, with symptoms of bloody diarrhoea, abdominal pain, and rectal bleeding (1, 2). Although the etiology is unknown, the pathogenesis is likely dependent on the interactions between local immunity and environmental factors in genetically susceptible individuals. Several animal models were developed to understand the pathology of IBD, but no specific model existed for either disease. Gut inflammation can be induced using chemical compounds e.g. DSS or by genetic targeting of specific genes e.g., regulatory cytokines. The majority of the knockout (KO) mice develop colitis with histopathology resembling UC (e.g., IL-2 KO mice) or entero-colitis (e.g., IL-10 KO mice) (3). Okayasu and colleagues described a model in which mice, receiving DSS orally, developed acute and chronic colitis resembling UC. The mechanism by which DSS induces intestinal inflammation is unclear but is likely the result of damage to the epithelial monolayer lining of the large intestine (4). The damage to the epithelium may lead to the dissemination of pro-inflammatory intestinal contents (e.g. bacteria and their products) into underlying tissue. The resulting inflammation has been shown to include polymorphonuclear cells, macrophages, and B and T cells. The mucosal inflammation is due to either an excessive Th1- or an excessive Th2-T cell response. Th1-response is usually characterized by the increased IL-12, IFN-γ, and TNF-α production while increased IL-10 and/or IL-4 production can be the signatures of Th2-response (5, 6). The DSS colitis model is very popular in research due to its rapidity, simplicity, reproducibility, and controllability. Acute, chronic and relapsing models of intestinal inflammation can be achieved by modifying the concentration of DSS and the frequency of administration (7). To develop robust colitis in mice, different dosage of DSS was used and the optimum DSS dose was found to be dependent on the mouse strain. The optimal concentration recommended, for inducing robust colitis over a period of 7 days, was 1.5% and 3.0% in C57BL/6 and 2.5-5.0% in BALB/c mice. To induce chronic colitis, mice were typically subjected to a low dose of three to five cycles of weekly DSS exposures, each dose was followed by a 1- or 2-week rest period (1, 3, 8–10). Till now there is no such model where all the stages of colitis can be studied in a single model. In addition, there is no common optimised dose of DSS with zero mortality rates that can cause acute colitis within the same time period for both strains of mice.

In the present study, we used C57BL/6- and BALB/c-mice to study the differential effects of a common dose of DSS. The uniqueness of the DSS model in the current report is its rapidness and compositeness. In this model, within the two weeks’ time period all the stages of colitis can be studied. The optimized dose is able to generate acute colitis within the same time period for both the mice strains. In the current report, the results revealed that the immunological background of the host plays critical role in the onset of colitis. Up regulation of the Th1 response, in T cells following DSS treatment, enhanced the severity of colitis more in C57BL/6 than in BALB/c mice. Th1 background of C57BL/6 mice might have led to develop the higher inflammatory outcome and chronicity of colitis, while Th2 background of BALB/c mice might have helped to alleviate the disease or complications at an earlier stage.

## Results

During treatment with DSS by oral administration, all mice of both the strains developed diarrhoea with significant weight loss, and rectal bleeding. They also turned positive in faecal occult blood test. C57BL/6 and BALB/c mice displayed similar physiological, clinical and histo-pathological changes during the acute phase of colitis when the DSS dose was high (5% for first 7 days of treatment). At a lower (2.5%) dose of DSS for next 7 days, acute symptoms turned into chronic in C57BL/6 mice while BALB/c mice directly moved to a recovery phase without going through a chronic phase. The severity and the rate of disease progression were much higher in C57BL/6 mice compared to BALB/c even in the acute phase of the disease. Details of the findings associated with these observations are elaborated successively in upcoming sections.

### Optimum DSS dose for rapid colitis induction-

Various dosages of DSS were orally administered in C57BL/6- and BALB/c-male mice. Untreated mice were used as time matched control in the study. To study the stages of the colitis within a shorter time scale (Fig.2A), both the mice strains were treated with 7.5% DSS for a period of 2 weeks. The dosage of 7.5% DSS was tolerated by BALB/c; it was lethal for C57BL/6. C57BL/6 mice started dying following 2 days of 7.5% DSS treatment. To overcome these problems, both strains of mice were treated with a lower dose of DSS i.e. 5% of DSS for 2 weeks. Outcome of the treatment with a lower dosage of DSS was same, i.e. 5% DSS for 2 weeks was lethal for C57BL/6 while BALB/c tolerated. We modified the treatment plan and now both the mice strains were treated with DSS at a concentration of 5% for the first week followed by a reduced dose of 2.5% DSS for the 2^nd^ week. At this newer dosage (5%+2.5%) of DSS, no mortality of either of the strains was observed. Results further revealed that at the dosage (5%+2.5%), the stages of the disease were prominent. Unless otherwise described, DSS dosage of 5%+2.5% was used in all further studies. Following standardization of DSS dosage, physiological changes of treated animals were monitored as the primary and most important sign of disease progression and compared it with the corresponding control groups.

**Figure 1:**
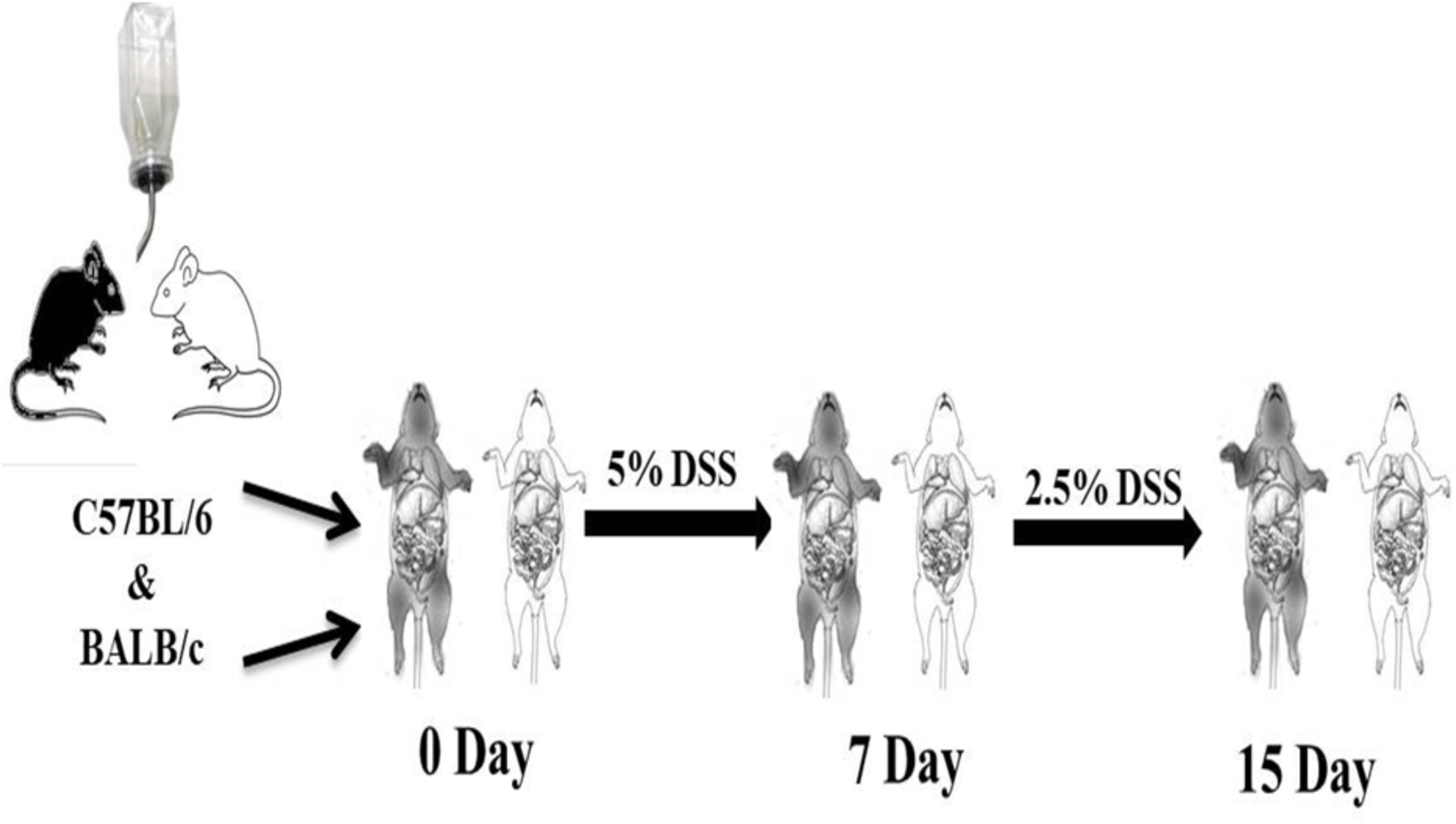
Schematic diagram to illustrate the induction of colitis in C57BL/6 and BALB/c mice. Combined dose of DSS to study acute, chronic and recovery phase of colitis within a period of two weeks with zero mortality rate. DSS dose was similar for both the strain of mice.

**Figure 2.**
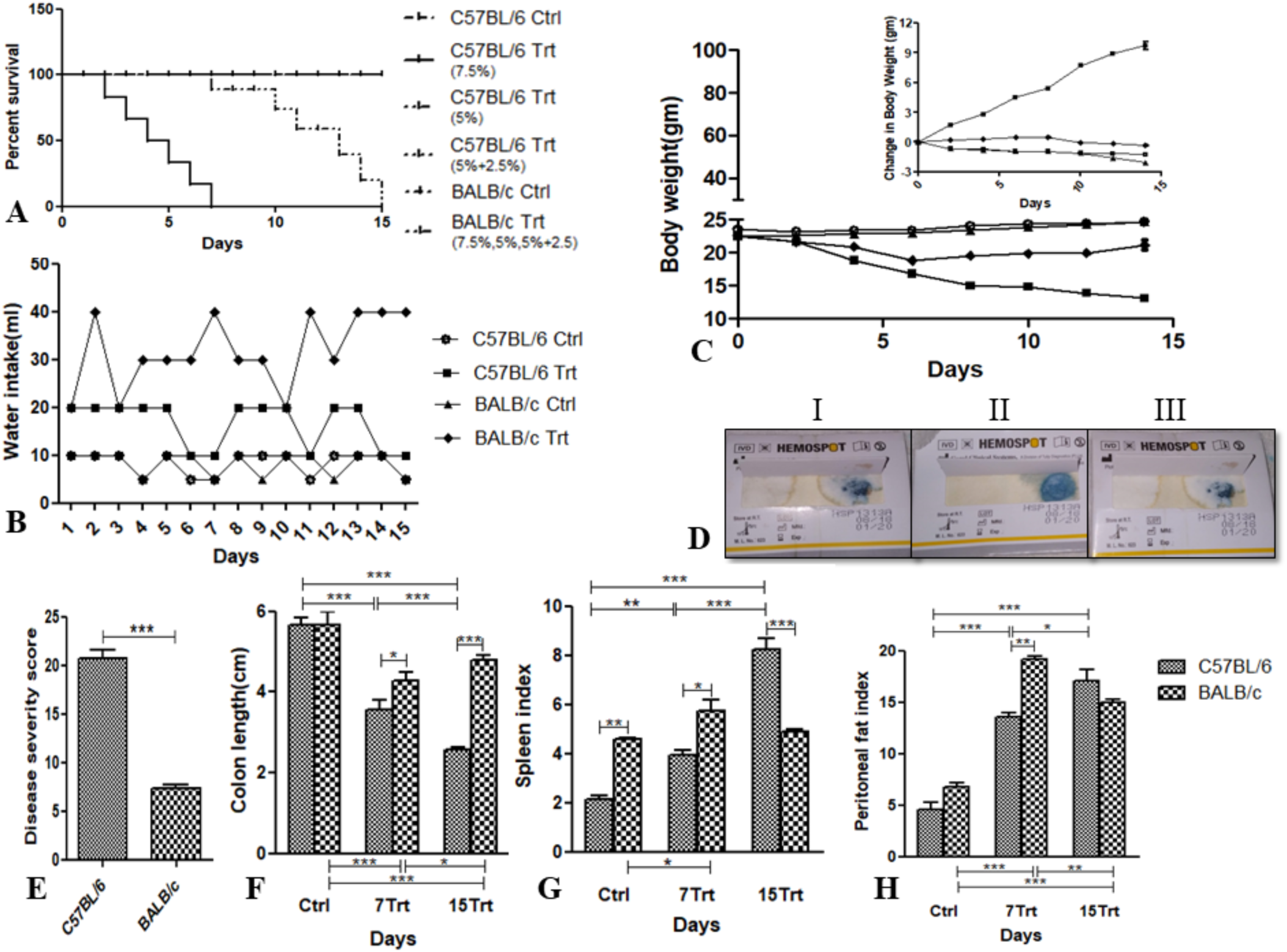
Survival rates of mice following challenges with different doses of DSS and its effect on physiology. (A) Male C57BL/6 and BALB/c mice were challenged with 7.5% or 5% or combined doses of 5%and2.5% DSS in their drinking water to find a suitable dose with no mortality at least for a period of two weeks. The Kaplan–Meier plot represents the percentage survival of control groups with its respective treatment groups (n=6 animals). Finally, mice were challenged with combined doses of 5% and 2.5% DSS for a period of two weeks, in the manner of 5% DSS for one week and then 2.5% for another one week for further studies, (B) Daily water intake of both control and DSS treated C57BL/6 and BALB/c mice for 15 days. Water intake of the treated groups was compared with the control groups, (C) Body weight of control and treated groups were measured every alternative day for up to 15 days in C57BL/6 and BALB/c mice. (D) Presence of faecal occult blood was measured from stool samples of C57BL/6 (D: II) and BALB/c (D: III) treated group of mice by Hemospot kit (standard guaiac method) and compared with the untreated group as well as absolute positive and negative control provided in the kit (D: I). The different intensities of blue color indicated various levels of occult blood in stool, e.g. no color meant no occult blood, a light blue meant around 5mg/dl occult blood, and strong blue implied more than 5mg/dl occult blood in stool, (E) Disease severity of both treated C57BL/6 and BALB/c mice were measured and compared based on stool consistency, amount of occult blood in stool and rectal bleeding, (F) Colon length in cm of control and DSS treated group of C57BL/6 and BALB/c were measured on days 0 (Ctrl), 7 (7Trt), and 15 (15Trt), (G) Spleen indices for control and treated groups were determined in C57BL/6 and BALB/c mice on days 0, 7, and 15. Spleen index was calculated by dividing spleen weight (isolated from euthanized mice) on specified day by body weight of that day, (H) Peritoneal fat indices for both control and treated groups were measured in C57BL/6 and BALB/c mice on days 0, 7, and 15. Peritoneal fat index was calculated by dividing peritoneal fat weight (isolated from euthanized mice) on specified day by body weight of that day. All data presented as the means ± SD (n=6). Two-way ANOVA followed by Bonferroni test was performed to determine the significance level for all the analysis. **\*** corresponds to P<0.05, **\*\*** corresponds to P<0.01, **\*\*\*** corresponds to P<0.001.

### Distinct changes in physiology at the different time point-

Oral administration of DSS (5%+2.5%) for a period of 2 weeks induced notable physiological changes in both mice strains. DSS intake was differed a lot between control and treated groups for both the strain of mice (Fig.2B). Control group of mice for both the strains were consumed nearly equal amount of water but water consumption of treated group for both the strains were significantly higher. BALB/c mice consumed more DSS containing water than C57BL/6 mice. Although DSS consumption was higher in treated BALB/c mice but the weight loss, diarrhoea, rectal bleeding and other physiological changes were more intense in treated C57BL/6 mice. During the period of DSS treatment, gradual weight loss was noticed among treated C57BL/6 mice throughout the treatment period (Fig.2C). For BALB/c weight loss was seen in the first week of DSS treatment. After that, as the DSS dose decreased, BALB/c mice were started gaining weight and reached to the normal condition at the end of the treatment (Fig.2C). Same trend was seen for diarrhoeal severity, rectal bleeding and presence of faecal occult blood (Fig.2D). Based on the above mentioned criteria disease severity score (Table.1) was calculated and found that disease severity score of treated C57BL/6 mice was significantly higher than treated BALB/c mice (Fig2E.). Colon length shortening, spleen index and peritoneal fat index were also measured to determine the severity of colitis. Consistent shortening of colon was seen in C57BL/6 but for BALB/c it was seen for up to 7days, then it was reached to almost normal level on 15^th^ day (Fig.2F). Blood loss and anaemia were the most common complications of colitis, which ultimately caused splenomegaly in host. In the current study spleen index was significantly higher in the treated group of mice (Fig.2G). Peritoneal fat index was also higher in treated group, to create a niche for inflammation (Fig.2H). Both the indexes were gradually increased for the whole treatment period in the treated C57BL/6 group, whereas both the indexes were becoming normal for treated BALB/c group after the decrease of DSS on the second week of treatment period. Distinct physiological changes of treated groups, especially shorter colon length and faecal occult positive test as well as rectal bleeding were the indication of distinct histo-pathological changes of colon in the treated groups compared to the control groups.

**Table 1:** Disease severity score based on stool texture and rectal bleeding

| Condition | Score |
| --- | --- |
| Normal + no hemocult | 0 |
| Soft + no hemocult | 1 |
| Soft + hemocult | 2 |
| Soft+ very little amount of rectal bleeding | 3 |
| Very soft+ blood | 4 |
| Watery+ rectal bleeding | 5 |

### Histo-pathological assessment of colitis based on colonic lesions-

Oral administration of 5% DSS for first 7 days showed almost similar amount of severity for both the strains. The histological changes of the colon at that particular time point was characterized by the parameters listed in Table.2 (12). Colon sections of control mice were (Fig.3A-B, 3G-H) showed the intact epithelium, well defined crypt length, and no edema neutrophil infiltration in mucosa and sub-mucosa, and any ulcers or erosions. In contrast, colon tissues from DSS treated mice were showed increasingly severe inflammatory lesions extensively throughout the mucosa during the first week of DSS treatment (Fig.3C-D, 3I-J). Ulcers, shortening and loss of crypts were seen in the whole colon. For C57BL/6 strain this condition was persisted up to the end of the treatment (Fig.3E-F). Even at the lower dose of DSS did not show any significant recovery. However lower dose of DSS for last 7 days helped BALB/c mice to recover from its previous histo-pathological condition. Clearing of neutrophils, lymphocyte from lamina propria was started as a part of the recovery process (Fig.3K-L). To measure the severity histo-pathological score was given (Table.2). This score was significantly higher for the treated C57BL/6 mice compare to the treated BALB/c mice (Fig.3M). Ulceration, edema, neutrophil infiltration in mucosa and sub-mucosa and severe inflammatory lesions and epithelial injury were thought to be the initial event of epithelial cell barrier function loss. To prove it further gut integrity was measured in treated and control group of mice.

**Table 2:**
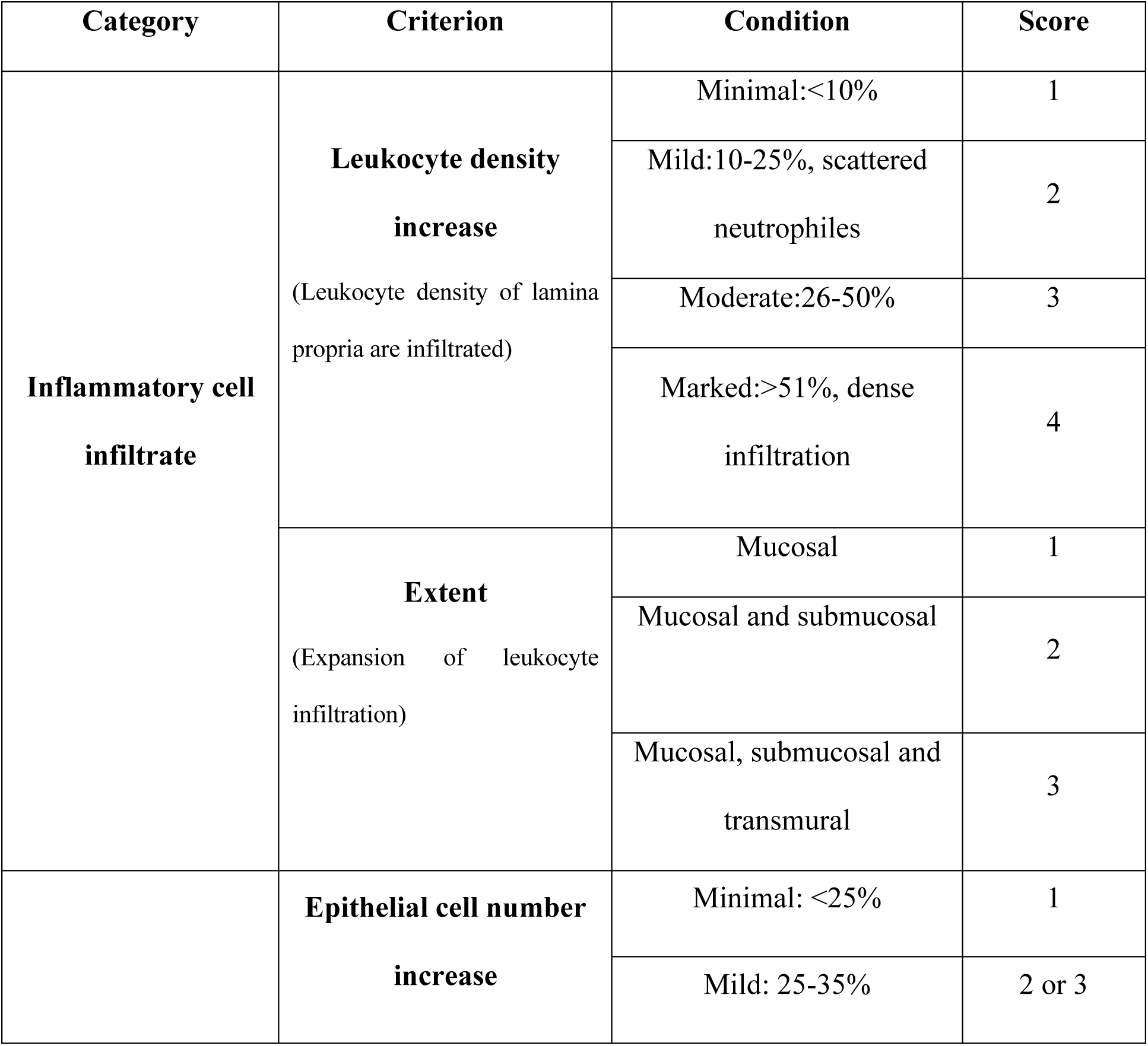
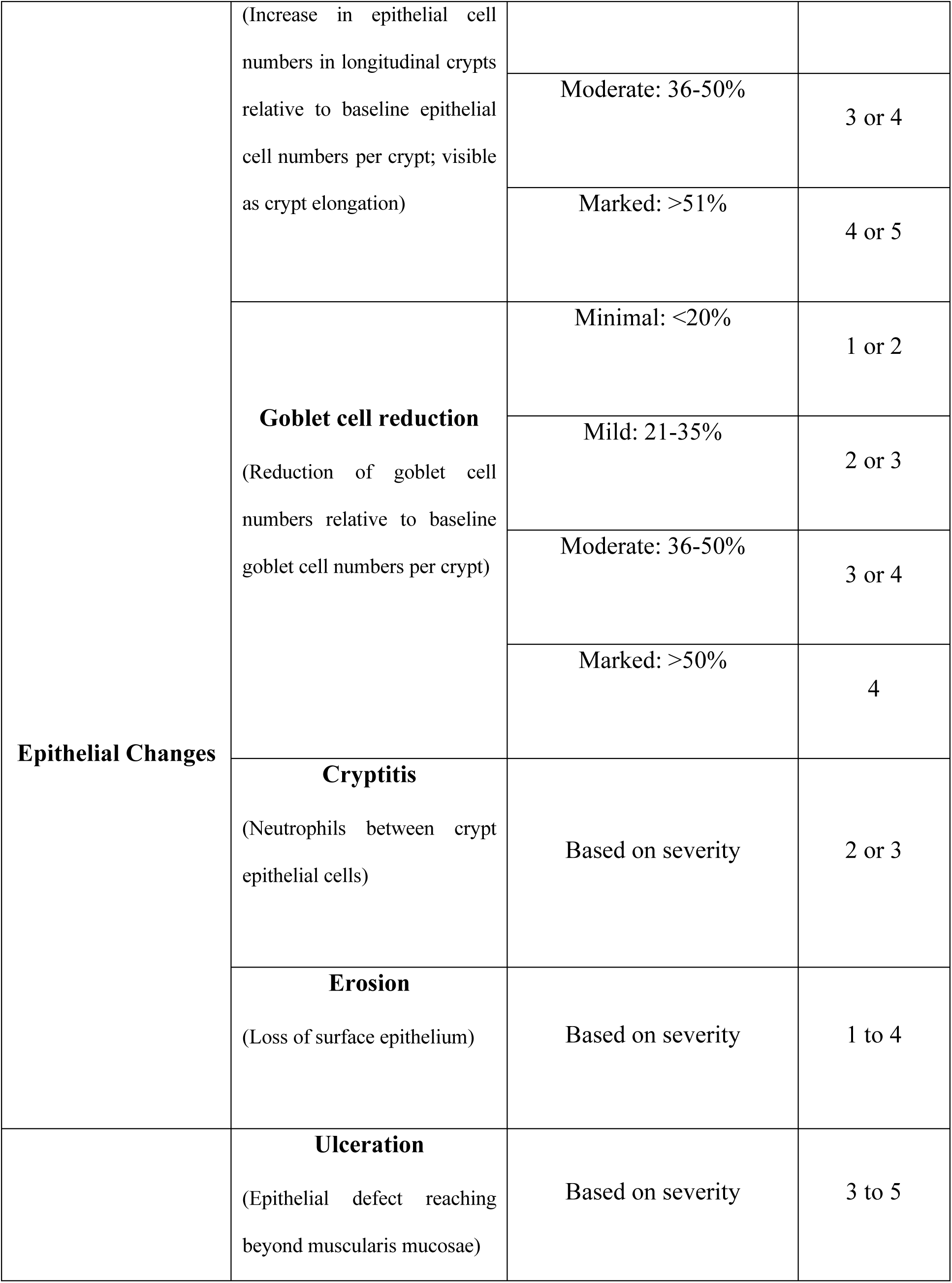
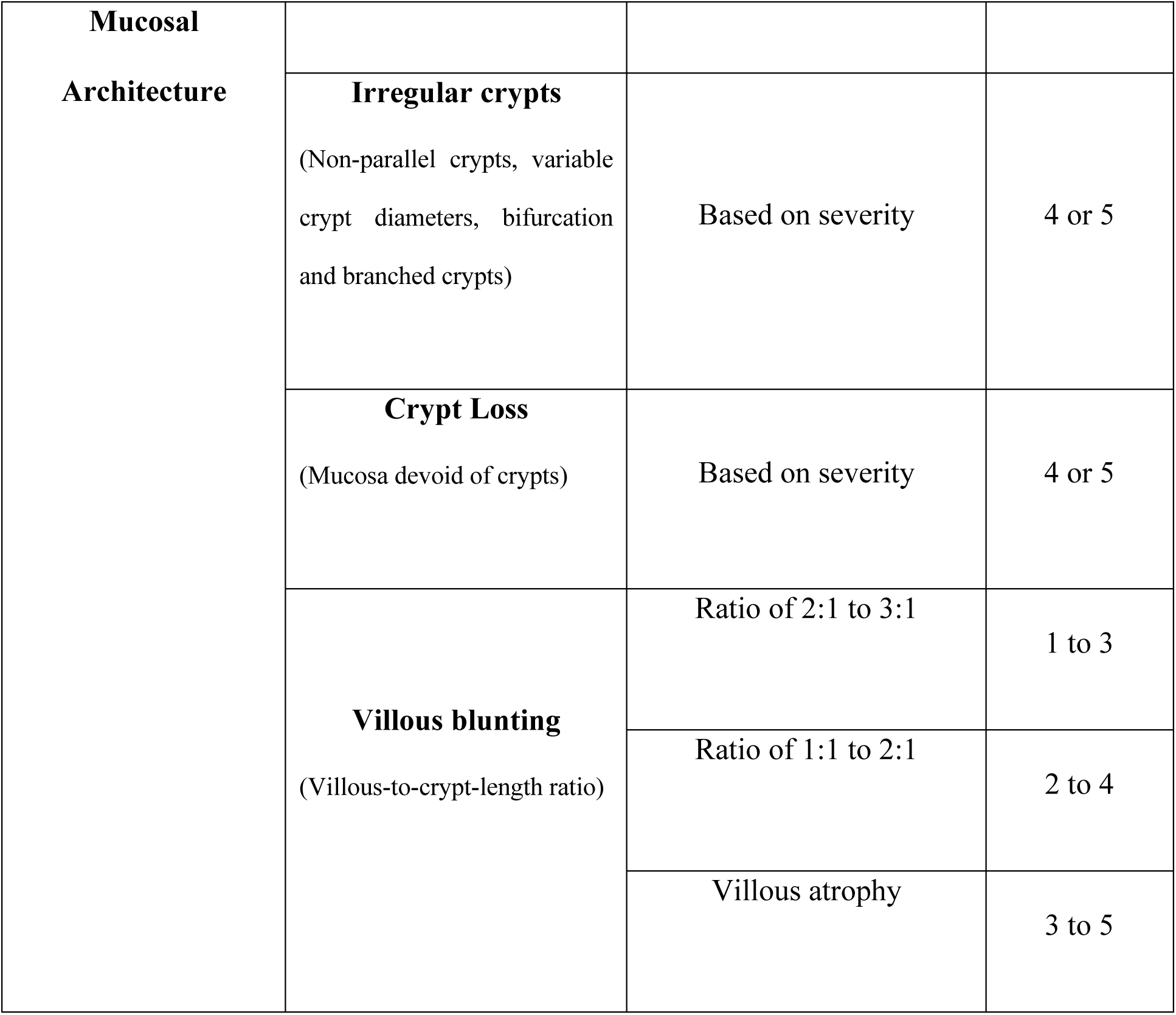
Scoring method used for Histo-pathological analysis

**Figure 3.**
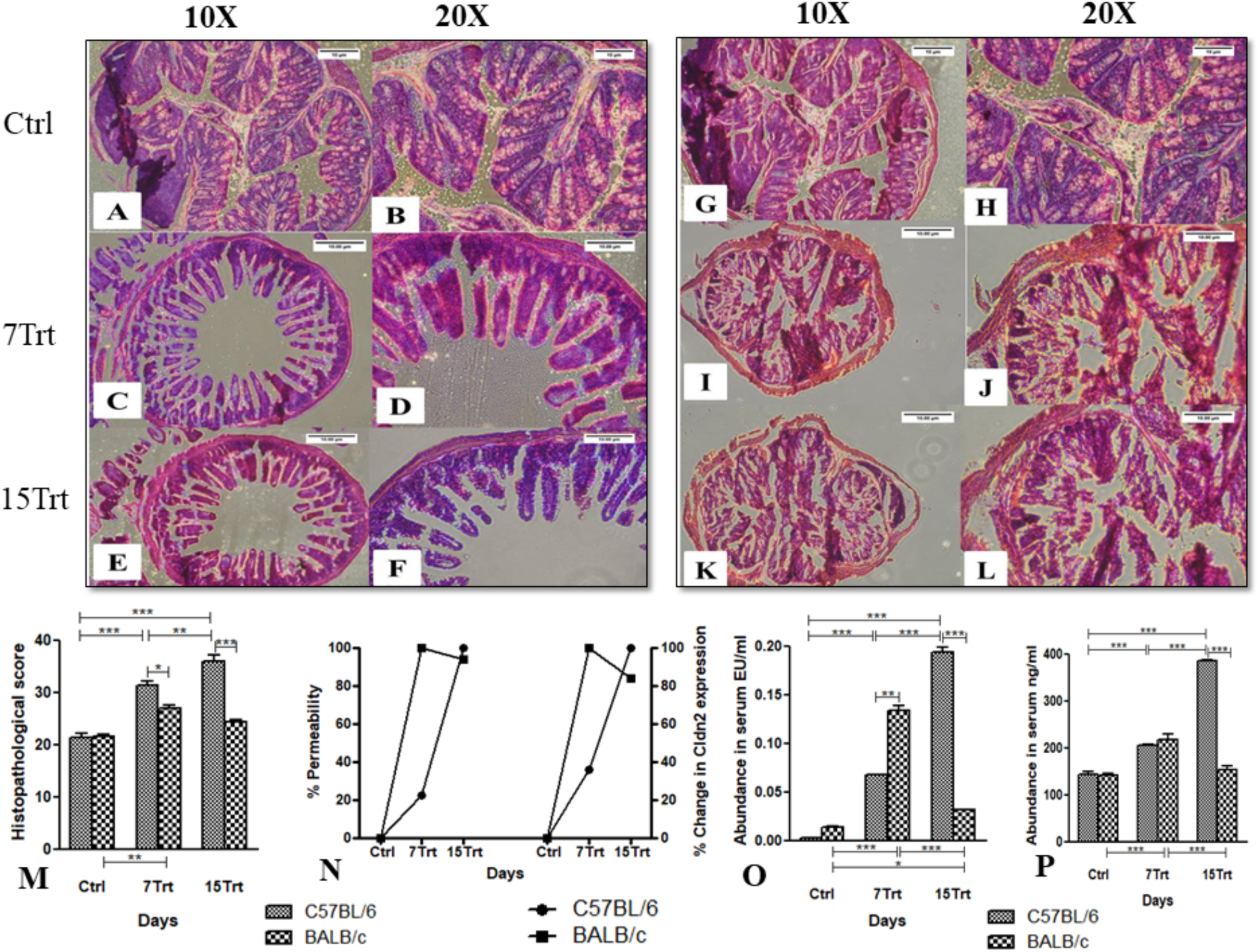
Changes in colon histopathology as well as intestinal barrier function following DSS challenge and its effect on innate defence mechanism. Colon sections were stained with Hematoxylin-eosin for control and DSS treated C57BL/6 (A-F) and BALB/c (G-L) mice on 0day (Ctrl), 7^th^ day (7Trt) and 15^th^ day (15Trt) to measure the changes in epithelium and mucosal architecture and inflammatory cell infiltration. (M) Histo-pathological scoring was done based on above mentioned criteria for C57BL/6 and BALB/c mice. (N) Levels of FITC-dextran in the serum of control and DSS treated C57BL/6 and BALB/c mice were measured as an indicator of intestinal permeability on 0day (Ctrl), 7^th^day (7Trt) and 15^th^ day (15Trt) and depicted as % permeability. Changes in mRNA expression level of tight junction gene, claudin2 (Cldn2) was measured from colon tissue by qRT-PCR for both control and treated group of C57BL/6 and BALB/c mice and depicted as % change in gene expression. (O) Endotoxin concentration in the serum was measured for both control and treated group of mice in C57BL/6 and BALB/c strain as a consequence of leaky gut. (P) Antimicrobial peptide, Lipocalin2 (LCN2) level was measured in serum of C57BL/6 and BALB/c mice on 0day (Ctrl), 7^th^ day (7Trt) and 15^th^ day (15Trt) as a marker of host defence mechanism against serum endotoxin. All data are presented as the means±SEM (n=6). Two-way ANOVA followed by Bonferroni test was performed to determine the significance level. **\*** corresponds to P<0.05, **\*\*** corresponds to P<0.01, **\*\*\*** corresponds to P<0.001. (For histological analysis, pictures were taken with magnification of 10X, scale bar 100µm and 20X, scale bar 50 µm).

### Compromised gut integrity and high Claudin2 mRNA expression in DSS treated mice-

Profound structural loss and inflammatory cell infiltration inside colon from the histopathology was an indication of compromised gut barrier function (23, 24). To prove it further, serum FITC-Dextran level was measured following oral administration of DSS. High concentration of FITC-Dextran in serum was the indication of high gut permeability. Serum concentration of FITC-Dextran was converted into the % permeability. Fold change of the FITC-Dextran concentration of treated groups were calculated with respect to the serum FITC-Dextran concentration of control which further converted in to % permeability for better understanding. In case of C57BL/6 % permeability was increased gradually in treated group compare to the control and reached its highest point on 15^th^ day of DSS treatment. For BALB/c the scenario was a bit different. Gut permeability was highest on 7^th^ day for the DSS treated group and then started decreasing gradually as the DSS dose was decreased (Fig.3N).

Tight junction proteins played a very crucial role in maintaining gut permeability. Some of their expressions were down regulated in leaky gut epithelia e.g. Claudin-1, Occludin-1, but some were overexpressed. Claudin-2, was one such tight junction protein, which was predominantly expressed in leaky epithelia, known as a channel-forming tight junction protein permeable to small cations and water (25). Earlier reports also showed that Claudin-2 was overexpressed in colitis patients (26, 27). Therefore, transcription level expression of claudin2 gene was measured from colon tissue as a potential marker for leaky gut. Fold change of claudin2 gene expression in treated mice of both the strain were converted in to percentage change of claudin-2 expression and compared with its respective control group. Claudin-2 expression was sharply increased in both C57BL/6 and BALB/c mice on the 7^th^ day of DSS treatment. Whereas on the 15^th^ day, when the DSS dose became low, gradual increase of claudin-2 gene expression was occurred for treated C57BL/6 group, but for treated group of BALB/c, gene expression was started decreasing and tends to the normal level. So the claudin-2 expression was highest in both the strain of mice, where there was highest gut permeability. (Fig.3N). At the diseased condition, compromised gut barrier function opened up the way for gut luminal microbes mostly pathogens to migrate to the host circulatory system, which ultimately caused systemic infection in host. Among the pathogenic microbe gram negative bacteria released endotoxin which promote the activation of host defence mechanism by activating the inflammatory responses of host.

### Presence of bacterial endotoxin in host serum as a consequence of leaky gut-

Enhanced intestinal permeability increased the risk of bacterial translocation from the gut lumen to the host circulatory system and also increased the chance of endotoxemia (28, 29). The serum level of endotoxin (LPS) was examined in both the mice strain for control and treated mice. The amount of endotoxin level in serum was highest at the point where the gut permeability was the highest. For C57BL/6 it was on 15^th^ day and for BALB/c on 7^th^ day (Fig.3O). Elevated endotoxin level in the host serum, activated host innate defence mechanism by producing different kind of antimicrobial peptides, which ultimately help the host to cope up from the diseased condition and maintain the homeostasis of the system.

### Host defence mechanism against bacterial endotoxin-

The innate immune system initiated host defence against invasive bacterial pathogens, bacterial endotoxin (LPS), and using specific recognition mechanisms. Antimicrobial peptide was one of the main and early host defence response, directly interact with LPS to inhibit the release of inflammatory cytokines and thus induced an anti-inflammatory effect (30, 31). Lipocalin2 (LCN2) was one of such antimicrobial innate immune glycoprotein which played a role in colitis by mitigating gut injury and maintaining iron homeostasis (32, 33). LCN2 level was measured from the serum of control and treated group of mice. The LCN2 amount was highest at the point where the highest amount of endotoxin was present in the serum. For C57BL/6, it was highest on 15^th^ day and for BALB/c on 7^th^ day (Fig.3P). Production of LCN2 was stimulated by the binding of toll like receptors mainly toll like receptor 2 and 4 of immune cells with the pathogen (34). To investigate the presence of high LCN2 in host system, mRNA expression level of toll like receptors 2 & 4 (*TLR* 2&4) were measured from colonic tissues of mice as an activator of LCN2. The data was supporting the serum LCN2 data. Serum LCN2 level was highest on 15^th^ day for C57BL/6 and 7^th^ day for BALB/c treated group. mRNA expression of *TLR2* and *TLR4* genes were also showed the same trend like LCN2.The expressions of both *TLR 2* and *4* was highest on15^th^day for C57BL/6 treated group and for BALB/c treated group it was on 7^th^ day and after 7^th^ day it was started decreasing to the normal level. (Fig.4A). High *TLR* expression was the indication of activation of pro-inflammatory immune response of host. To know the inflammatory responses of treated animals, expression of inflammatory cytokines were measured as a consequence of higher expression of antimicrobial peptide and specific toll like receptors.

### DSS induced the production of inflammatory mediators-

The results so far revealed a) enhanced intestinal permeability and b) accompanying immune cell infiltration in the colon of the either type of mice. Enhanced permeability and immune cell infiltration could be the main reason to increase mucosal production of pro-inflammatory cytokines. Pro-inflammatory cytokines might play a pathogenic role in colitis due to DSS treatment in C57BL/6 and BALB/c mice. To estimate the extent of inflammation during colitis development, transcriptional profiling of selected cytokines and Myeloperoxidase (MPO) in colonic tissue homogenates was measured. The chosen genes, i.e., *TNFα, IFNγ, IL1β, IL6, IL12, IL17, IL21, IL10*, and *MPO*, had been implicated in the pathogenesis of colitis (Fig.4B-D). At the higher dose of DSS (5%) activation of Th1 and Th17 inflammatory response were quite similar in both the strain of treated mice. On first week of treatment, all the Th1 and Th17 inflammatory cytokines (e.g. *TNFα, IFNγ, IL1β, IL6, IL12, IL17, IL21*) (9) and MPO gene expression were up regulated. Although mRNA expression of all those genes were significantly higher on 7^th^ day (following treatment with 5% DSS) in both the treated strain but compare to BALB/c treated mice, all the gene expressions were significantly higher in C57BL/6 treated group. Transcriptional changes in *IL10* cytokine level was measured as an activator of Th2 inflammatory response (9). Expression of *IL10* was significantly down regulated in both the strain of treated mice on 7^th^ day (following treatment with 5% DSS), when the activator of Th1 and Th17 inflammatory cytokines were significantly up regulated. On the second week when the DSS dose was low (i.e.2.5%) expression of Th1 and Th17 cytokines were again significantly up-regulated in treated C57BL/6 mice on 15^th^ day (treated with 2.5% DSS) compare to its 7^th^ day (treated with 5% DSS). No significant up regulation was observed in the Th1 and Th17 cytokine levels on 15^th^ day (followed by 2.5% DSS treatment) compared to 7^th^ day (followed by 5%) in DSS treated BALB/c mice. Most of the gene expressions were even significantly down-regulated and became normal on 15^th^ day (when the DSS dose was lower i.e.2.5%) compare to the 7^th^ day (when the DSS dose was higher i.e.5%) in treated BALB/c mice. As the Th1 and Th17 cytokines were significantly up regulated throughout the treatment period in C57BL/6 treated group, that was might be the reason of no significant up-regulation of Th2 cytokines 15^th^day (treated with 2.5% DSS) compare to 7^th^ day (treated with 5% DSS). But for BALB/c treated group Th2 cytokine level was significantly up-regulated on 15^th^ day (treated with 2.5% DSS) compare to 7^th^ day (treated with 5% DSS). Normal or prone to be normal Th1 and Th17 cytokine levels were might be the reason behind it. High mRNA expression of inflammatory markers of treated mice compare to the control was the indication of activation of inflammatory responses of host. To confirm it further, protein level expression of inflammatory markers was measured in both control and treated group of mice.

### Colonic MPO Activity-

mRNA expression of MPO gene was significantly higher in DSS-treated groups than corresponding controls for all the period of study. To prove it further MPO level was measured in protein level. It was mainly a heme-containing peroxidase expressed mainly in neutrophils and to a lesser degree in monocytes. In the presence of hydrogen peroxide and halides, MPO catalysed the formation of reactive oxygen intermediates, including hypochlorous acid (HOCl). The MPO/HOCl system played an important role in microbial killing by neutrophils (35). Colonic MPO activity was measured as an indicator of the extent of neutrophil infiltration into the mucosa (36, 37). Fig. 4E showed the quantification in concentrations of this parameter in the experimental and control animals. It was found that MPO activity reached to the maximum level on 15th day for C57BL/6. For BALB/c MPO activity was more or less similar on 7^th^ day and 15^th^ day.

**Figure 4.**
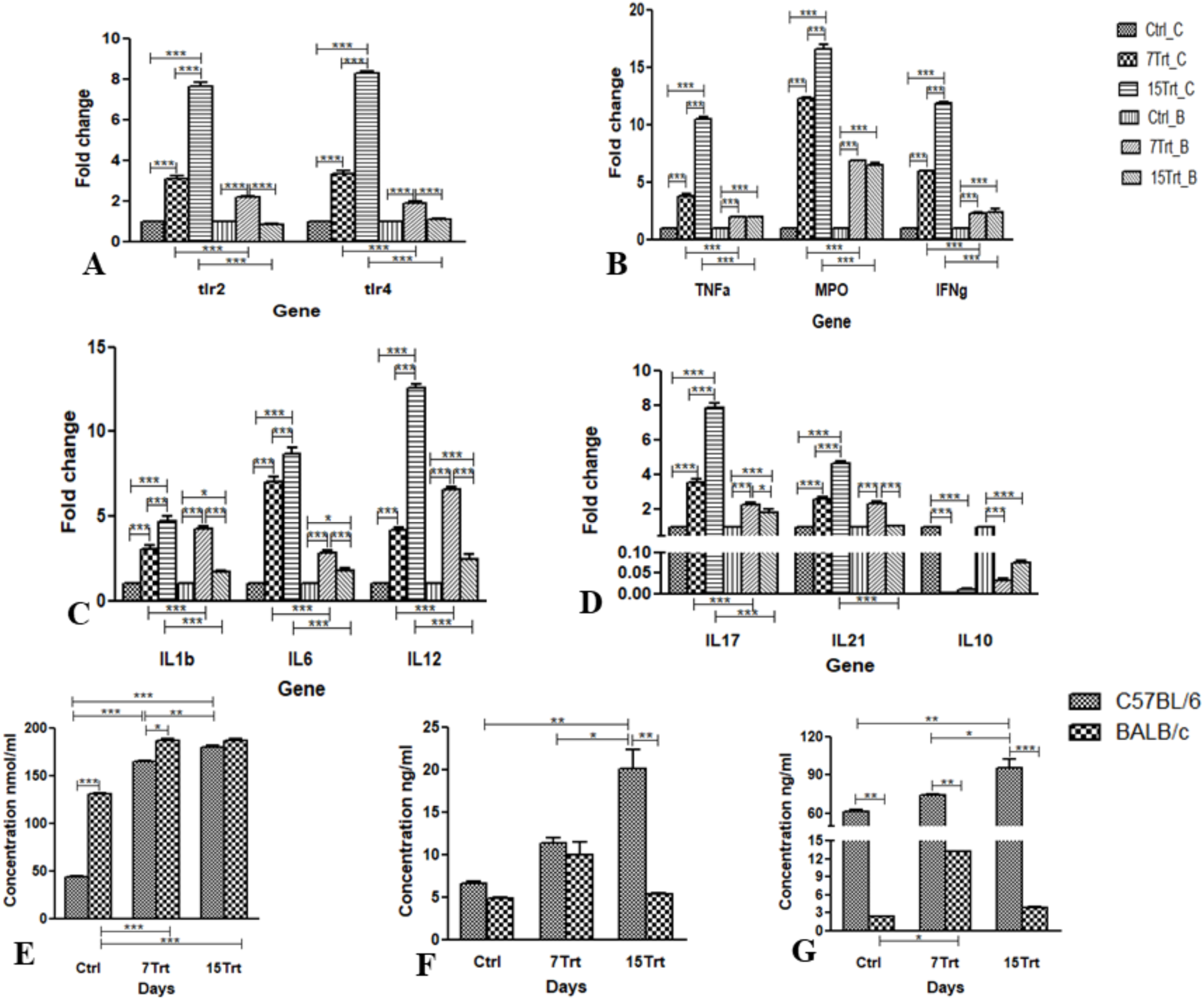
Transcriptional and protein level gene expression of inflammatory markers in colon tissue and cecal content after DSS treatment. (A, B, C, D) Kinetics of toll like receptors and pro and anti-inflammatory markers level were measured in both C57BL/6 and BALB/c by qRT-PCR. Statistical significance was calculated by two-way ANOVA followed by Bonferroni test. **\*** corresponds to P<0.05, **\*\*** corresponds to P<0.01, **\*\*\*** corresponds to P<0.001. All data are depicted as means±SEM values of 3 biological replicates. (E) MPO enzymatic activity as an index of neutrophils infiltration was measured in colon tissue of C57BL/6 and BALB/c mice on 0day (Ctrl), 7^th^ day (7Trt) and 15^th^ day (15Trt). (F) C-reactive protein, level was measured in colon tissue of C57BL/6 and BALB/c mice on 0day (Ctrl), 7^th^ day (7Trt) and 15^th^ day (15Trt) to know the acute phase response of host due to inflammation. (G) Cecal secretory IgA level was measured as an up-regulated immune response of host in C57BL/6 and BALB/c mice on 0day (Ctrl), 7^th^ day (7Trt) and 15^th^ day (15Trt). Data are presented as means±SEM (n=6). Two-way ANOVA followed by Bonferroni test was performed to determine the significance level. **\*** corresponds to P<0.05, **\*\*** corresponds to P<0.01, **\*\*\*** corresponds to P<0.001.

### C- reactive protein (CRP) production from colon-

C-reactive protein, mainly known as acute phase response protein, showed flare of inflammatory reactions to activate host innate immune system in diseased condition. It was an ancient highly conserved molecule and a member of the pentraxin family of proteins. The function of CRP was felt to be related to its role in the innate immune system. It activated complement system, binds to Fc receptors and acted as an opsonin for various pathogens. Interaction of CRP with Fc receptors lead to the generation of pro-inflammatory cytokines that enhance the inflammatory response (38). The synthesis of CRP was inducible by TNFα and IL6 (39). It was found that colonic TNFα level was high in the colon of DSS treated mice as a cause of inflammation. Colonic CRP level was measured by ELISA as an effect of high TNFα mRNA expression in colon (Fig.4F). Colonic CRP became significantly high on 15^th^ day for C57BL/6 strain compare to the control. Although there was a significant change in TNFα mRNA expression in colon for BALB/c strain, but no significant change was found in colonic CRP level.

### Microbial IgA Production-

Immunoglobulin A (IgA), as the principal antibody class in the secretions that acted as an important first line of defence. The vast surfaces of the gastrointestinal tract represented as major sites of potential attack by invading micro-organisms. Fecal/cecal IgA evaluate immunological respond to intestinal pathogens (40). Th17 cytokine promoted IgA secretion in colonic mucosa (41). From the transcriptional analysis it was clear that colonic expression of Th17 cytokine was higher in treated mice compare to the control. Same trend was found for IgA protein analysis from cecal content (Fig.4g). Like Th17 mRNA expression, the highest IgA produced by gut microbiota on 15^th^ day of DSS treatment for C57BL/6. For BALB/c it was on 7^th^ day.

## Discussion

DSS induced colitis, despite its shortcomings, has been of great use in understanding the pathophysiology of intestinal inflammation. This model, in skilled hands, represents a powerful tool to study a) the contribution of various aspects of the disease and b) to evaluate interventions designed to prevent or ameliorate disease. The DSS-induced colitis model will continue to yield valuable mechanistic clues regarding the interactions between host genetics, gut innate immunity, microbiota, diet and other environmental factors in maintaining gastrointestinal homeostasis.

In the present study a combined dosage of DSS was given for two weeks to study the differential effect of acute and chronic colitis as well as its recovery phase in two different immunologically biased mouse models. To the best of our knowledge, the current report is the first study where a common dosage of DSS was used to understand all the stages of colitis in two different immune biased mice model within a very short time period i.e. two weeks. Generally chronic symptoms persisted for months or longer and getting worse day by day whereas acute symptoms typically lasted for 7 to 14 days. Generally higher dosage of DSS (3-10%) was used to develop acute symptoms within 3-7 days and lower dosage (1-3%) for a prolonged time period minimum one week to several months was used to generate chronic symptoms (1, 3, 8, 42). To study the onset of acute colitis, 5% DSS was administered for the first 7 days in drinking water of C57BL/6 (Th1 biased) and BALB/c (Th2 biased). In acute phase the extent of colonic inflammation and other physiological changes were almost similar for both the strain of mice. Whereas the DSS dose become lesser i.e. 2.5% for the next 7days, to study the chronic symptoms, inflammatory condition of C57BL/6 become worst compare to the first 7days, but BALB/c started recovering from the inflammatory condition and trying to reach to the normal healthy condition. Although at the time of acute inflammatory condition Th1 and Th2 immune biased mice were responded in a similar manner, but for chronic condition their differential immunological background might play immense role to response them differently.

This disease classification was done based on clinical symptoms, molecular and histo-pathological changes (1, 11, 12). The long persisting inflammation, even after lowering the dose of DSS for C57BL/6 mice mimicked the chronic colitis condition. Whereas for BALB/c severe inflammation was persisted only for 7days in presence of higher DSS dose, which mimicked acute disease condition. In presence of lower dose of DSS, gradual decrease of inflammatory condition of BALB/c mimicked recovery phase of colitis (3, 7).

The clinical and histological changes were determined based on the phenotypic and pathologic changes such as stool consistency, diarrhoea, rectal bleeding, presence of faecal occult blood, body weight loss, colon shortening, spleen index and peritoneal fat index. These were the common phenomenon seen in DSS-induced experimental colitis (1, 11). Diarrhoea was due to the increased permeability of intestinal cells or hyper osmolarity in lumen led by DSS (4). Weight loss, shortening of the colon and high peritoneal fat index, as indicators for the severity of intestinal inflammation, high spleen index as indicator for anaemia and blood loss through rectal and faecal bleeding correlate with the pathologic and histological changes and were consistent markers for colitis (1, 4, 11). The data from current study also corroborated with the mentioned marker of colitis in the treated group of both the strain of mice.

Histological changes of colon were considered as one of the most important and consistent marker of colitis. Pathophysiological changes of treated group of mice compare to the control were a good indication to proceed further for histological analysis of colon tissue to determine the extent of inflammation. Acute histological changes in colon were determined by severe inflammatory lesions extensively throughout the mucosa, ulcers, shortening and loss of crypts in the whole colon and persisted for 7-10days. In chronic phase the same conditions were persisted for more than 7-10 days. Clearing of neutrophils, lymphocyte from lamina propria were seen as a sign of recovery period.

Diarrhoea, colonic ulceration, loss of crypts, infiltration of immune cells in to the lamina propria thought to be the initial inciting event that underlies compromised gut barrier function (23, 43). In the current study, following administration of FITC-Dextran orally, presence of high amount of FITC-Dextran in serum for both the strain of treated mice was the confirmatory test for high gut epithelial permeability. Gut permeability was highest on 15^th^ day in treated C57BL/6 group i.e. at the time of chronic condition. But for BALB/c highest gut permeability was observed at the acute phase i.e. on 7^th^ day.

Increased gut epithelial permeability was the indication of malfunctioned tight junctional complex. Generally in the healthy condition gut epithelial permeability depended on the apical plasma membranes of enterocytes and the junctional complexes between them. The main diffusion barrier within this junctional complex was formed by the tight junction, which contains trans-membrane proteins. Claudin-2, was one of such tight junction protein, which was predominantly expressed in leaky epithelia, was known as a channel-forming tight junction protein permeable to small cations and water (25–27). Claudin-2 mRNA expression was high for treated animals in acute and chronic condition and tends to normal at the recovery stage, same as the FITC Dextran data.

Enhanced intestinal permeability increased the risk of bacterial translocation to the host circulatory system and cause endotoxemia (presence of endotoxin in the blood) (28, 34). In the current study, at the treated condition when the gut permeability was high, serum endotoxin level became increased in chronic and acute phase as a consequence of leaky gut, and it was decreased in the recovery phase of BALB/c when the gut permeability was also less compared to the acute condition.

To manage this substantial exposure of endotoxin secreted by gram negative bacteria in to serum gut epithelial cells produced a diverse arsenal of antimicrobial proteins that directly killed or inhibited the growth of microorganisms (30, 44). Lipocalin2 was one of such antimicrobial peptide highly expressed in colitis patients, which binds with the ferric-siderophore complex of bacteria to inhibit bacterial growth (45, 46). In the present study both in acute and chronic condition Lipocalin2 expression was significantly high to pull down the serum endotoxin by killing pathogenic bacteria. But at the recovery phase it was trying to reach to the normal condition to balance the homeostasis.

Not only the activation of antimicrobial peptide production, endotoxin also formed complexes with lipopolysaccharide-binding protein and activate monocytes and macrophages through toll-like receptors(mainly *TLR2* and *TLR4*), involving the nucleotide-binding oligomerization domain 2 protein (NOD2) pathway (33, 46). In our study, as a result of high endotoxin in serum, mRNA expression of toll like receptor 2 & 4 started increasing in DSS treated mice at the acute and chronic condition of the disease, which further triggered the activation of pro-inflammatory cytokine e.g. TNFα, IL6 production in colon tissue.

TNF-α, produced by innate immune cells as macrophages, monocytes, and also by differentiated T cells and triggers Th1 cell differentiation. TNF-α exerted its pro-inflammatory effects through increased production of IL-1β and IL-6, expression of adhesion molecules, proliferation of fibroblasts and pro-coagulant factors, as well as initiation of cytotoxic, apoptotic, acute-phase responses, and inhibition of apoptosis. TNF-α, also stimulate IFN-γ secretion by binding to the death receptor 3 (DR3) (5). IFN-γ secretion was also mediated by IL12 and IL21, as a result of Th1 mediated inflammatory response(47, 48). Another T cell subset that played major role in inflammation was Th17, characterized by the production of IL17 cytokine. IL-17 in general induced the recruitment of immune cells to peripheral tissues, a response that required NF-κB activation after IL-17 receptor engagement. IL-17 also lead to the induction of many pro-inflammatory factors, including TNF-α, IL-6, and IL-1β, suggesting an important role for IL-17 in localizing and amplifying inflammation (49). Furthermore, TNF-α and IL-6, which were both produced by Th17 cells, not only support Th17 cell development but also synergized with IL-17 to enhance the production of pro-inflammatory mediators. IL10 secreted by CD4+ Th2 cells, triggers the Th2 cell differentiation in host (50).

In the current study it was found that transcriptional expression of all Th1 and Th17 biased cytokines e.g. *TNF-α, IFN-γ, IL6, IL1 β, IL12, IL21, IL17* were significantly up regulated and Th2 biased cytokine e.g. *IL10* was significantly down regulated in the colonic tissue of DSS treated mice on the first week of DSS treatment when the dose was high (5%). More interestingly the Th1 and Th17 responses were significantly higher in C57BL/6 mice compare to BALB/c. Th1 biased genetic background of C57BL/6 may played a role over here for more activation of Th1 biased inflammatory cytokines, whereas the Th2 biased genetic background of BALB/c was responsible for restricted activation of the same. Th1 biased immunological condition of C57BL/6 might be responsible for disease chronicity and showed the continuous increase of-Th1 biased inflammatory cytokines even at the lower dose of DSS.

For BALB/c the case was something different might be for its Th2 biased immunological condition, where the activation of Th1 biased inflammatory cytokines were seen at the acute phase of disease. In presence of a lower dose of DSS (2.5%) on the second week, most of the Th1 cytokines were started coming to its normal level.

Activation nature of Th2 biased cytokine *IL10* was totally different from Th1; it showed total opposite kind of respond of Th1 cytokine in both the strain of treated group of mice. mRNA expression of *IL10* was down regulated in treated C57BL/6 mice for the whole treatment period but for BALB/c treated group it down regulated only in higher dose of DSS mainly at the time of acute phase, followed by the up-regulation of its expression and tends to normal level at the time of recovery phase. Different immunological genetic background might be the reason behind these surprising results.

The inflammatory cytokines also played a major role in attracting neutrophils at the site of inflammation. The granules of neutrophil granulocytes contain a number of enzymes, most importantly myeloperoxidase (MPO), which were important in the combat against bacteria. MPO used as potential biomarker to study the disease progression of colitis (11, 35, 51). For this study, MPO expression was checked in colon both transcriptionally and as an enzymatic assay. In both the cases MPO expression was significantly high in DSS treated mice compare to the control.

Activation of acute phase response protein at the site of inflammation was also influenced by high pro-inflammatory cytokine at the inflammatory site. C-reactive protein was on of such acute phase response protein, produced at the site of inflammation in presence of high TNFα and IL6 cytokines (39, 52). For this study as a consequence of high TNFα and IL6 cytokines production in colon tissue CRP level was measured from colon tissue. TNFα and IL6 cytokine expression was might be not enough to induce the significant production of CRP on 7day for C57BL/6 compare to the control, but significant change was observed on 15^th^ day. Whereas for BALB/c there was no significant change in CRP level of treated mice compare to the control throughout the study period because of low expression of TNFα and IL6.

Th17 cell derived IL17 induced the production of IgA in the intestine. Segmented filamentous bacteria had been identified as a potent inducer of Th17 cells in the intestine, which helped in high IgA production. This IgA segregated the microbiota from attaching to the intestinal epithelium, and limiting contact with the intestinal immune cells (41). In present study significantly high IgA level was observed in the colonic content of DSS treated mice in acute and chronic phase of disease. IgA level became normal in the recovery phase of the disease. Elevated IgA level in the diseased condition may be correlated with the increased gut permeability. This elevated passage of bacterial antigens in the intestinal mucosa might lead to a concomitant increase in the immune response through specific antibody production for compensating this failure in the barrier function of the gut epithelium (53).

## Conclusion

In conclusion, oral administration of the particular dose of DSS i.e. 5% for one week then 2.5% and for another one week causes a reproducible colitis in both C57BL/6 and BALB/c mice. DSS induced colitis activates the Th1 and Th17 immune response in host system. ThusTh1 genetic background of C57BL/6 mice forced to show immense severity and chronic response in DSS treatment. Severity was gradually increased with time even after lowering the dose of DSS. Th2 genetic background of BALB/c, helped it to recover from the acute diseased condition after lowering the DSS dose. Disease severity of BALB/c mice was also significantly lesser than C57BL/6. The uniqueness of this model, it is rapid, reproducible. Within a very short time period one can study different stages of colitis and its related physiological and immunological condition as well as complications from a single model system. The current model is novel and more versatile and robust to understand the disease etiology.

## Experimental procedures

### Mice-

Specific Pathogen free 6-8 weeks old, male C57BL/6-or BALB/c-mice with body weight in the range of 18-22 g, were co-housed in poly-sulfone cage using corncob as bedding material. Mice of same strains were housed together. All animals were housed in a pathogen-free environment with a 12h: 12h light-dark cycle at temperature 24 ± 3° with the humidity around 55%. Standard pelleted food and autoclaved water were provided ad libitum. Committee for the Purpose of Control and Supervision of Experiments on Animals, Govt. of India (CPCSEA) approved this study and all experiments were done as per the approved guidelines.

### Colitis Induction and Sample Collection-

Dextran Sulfate Sodium (DSS, M.W.-50kDa, Fisher Scientific) was added to autoclaved drinking water (*ad libitum*) at a concentration of 5% for first one week. Dosage of DSS was reduced to 2.5% for the 2^nd^ week. Fresh DSS solution was prepared thrice in a week. Control group was fed with normal autoclaved water. In a separate study, we checked that the minimum dose required to develop colitis like symptoms in BALB/c was 2.5%-3% with a prolong treatment time. The dose of DSS at 2.5-3% was, however, lethal for C57BL/6 (Data not shown). To overcome this problem, both the strains were treated separately with 7.5% and 5% DSS for a period of two weeks and found that these higher doses was also lethal for C57BL/6. We, therefore, modified our dosage plan and selected 5%+2.5% dosage as described elsewhere for two weeks treatment period, i.e. 5% DSS for the first one week followed by 2.5% for the 2^nd^ week.

Mice were divided into 3 groups with 6 mice per group. Except the control (untreated) group other two groups (treated) received DSS water as described in Fig.1. Healthy control animals received only autoclaved water. Mice were sacrificed on 0-, 7- and 15-day. Samples from cecum, colon, blood, spleen, and peritoneal fat were collected on each of the days. Snap frozen tissue samples for protein analysis and cecum contents were stored at -80°C. Tissue samples used for RNA analysis were collected in RNA Later for further analysis. For histological studies tissue samples were stored in 4% PFA (paraformaldehyde) at the room temperature. Serum sample was prepared from whole blood by centrifuging it at 1600 rcf at 4°C for 10 minutes and stored at -20°C until further analysis.

### Measurement of Disease Severity: Symptoms and Inflammatory Score-

The parameters used, to assign the clinical symptoms to the DSS-treated and control mice, were water intake, body weight, stool consistency, occult blood in stool, rectal bleeding, and general health condition over the days of the treatment (1, 11). Based on the stool consistency, extent of occult blood in stool and rectal bleeding, severity level was scored separately and independently on a scale of 0-5 (Table1). At necropsy, colon length, spleen index and peritoneal fat index were measured in both control and treated group of mice.

### Histology-

Tissue sections (5µm thick) of mouse colon were stained with hematoxylin-eosin (H&E) to address the degree of inflammation. The stained tissues were analysed independently and in a blinded fashion with a standard inverted microscope (Olympus CKX53, Japan). The pathophysiology of the tissue was characterized by all of the following parameters, such as, the presence of ulcerations, inflammatory cells, signs of edema, loss of crypt epithelium, reduction in the number of goblet cell, and villous blunting (12). The parameters were scored on a scale of 1-5 and listed in the Table2.

### Intestinal Permeability Assay Using FITC-Dextran-

Control and treated mice of both the strain were food and water-starved for overnight and kept in a cage without bedding to limit the coprophagic behaviour (13). On the next day morning, body weights of mice were measured. FITC-dextran (Cat#F7250, Sigma-Aldrich, Missouri, US) dissolved in PBS (100mg/ml) was administered to each mouse (44 mg/100 g body weight) by oral gavaging. Following 4h of treatment, mice were anesthetised by isoflurane inhalation and blood was collected using a 1ml syringe with 25G needle by cardiac puncture. Serum was isolated from the blood using the protocol described previously and diluted with an equal volume of PBS. A volume of 100µl of diluted serum was added in each well to a 96 well micro plate in duplicate and fluorescence was measured with an excitation of 485 nm (20 nm band width) and emission of 528 nm (20 nm band width) (14, 15).

### RNA Extraction-

Total RNA was extracted from colon tissues using Qiagen RNeasy mini kit (Cat# 74104, Qiagen, India) according to the manufacturer’s protocol (16). In brief, 20-25 mg of frozen colon tissue in liquid nitrogen was homogenized by pestle-mortar and extracted in RLT buffer. Total homogenate in RLT buffer was centrifuged and transferred to RNeasy mini column and then washed using RW1 and RPE buffer. Total RNA was eluted in 30µl nuclease free water. Quality and quantity of extracted RNA was determined using NanoDrop 2000 machine (ThermoFisher Scientific, Columbus, OH, USA)by measuring ODs at 230, 260, and 280nm.

### cDNA Preparation-

mRNA in total RNA was converted to cDNAby using AffinityScript One-Step RT-PCR Kit (Cat# 600559, Agilent, Santa Clara, CA, USA) following the method described elsewhere (17). In brief, total RNA was mixed with random nonamer primer, Taq polymerase, and NT buffer. The mixture was kept at 45°C for 30 min for the synthesis of cDNA. To stop the reaction by deactivating the enzyme, temperature was increased to 95°C.

### Quantitative Real-Time PCR(qRT PCR)-

qRT-PCR reaction was set in a 96 well PCR plate using 30 ng of cDNA as template in the presence of 1μM/μl of each of forward (F) and reverse (R) primers (Table.3) for genes mentioned in Table3, SYBR green master mix (Cat#A6002, Promega, Madison, WI, USA), and nuclease free water. qRT-PCR was performed in Quantstudio 7 (Thermo Fisher Scientific, Columbus, OH, USA) using the cycle described as follows; 2 minutes at 92°C to activate DNA polymerase for 1 cycle, 15 seconds at 92°C for melting and 1 minute at 60°C for primer annealing along with extension of the chain and detection of the fluorescence for 40 cycles, then a program to find out the melting temperature of each product. All values were normalized with cycle threshold (C_t_) value of GAPDH (internal control) and fold change of the desired gene was calculated with respect to the control C_t_-value.

**Table 3:**
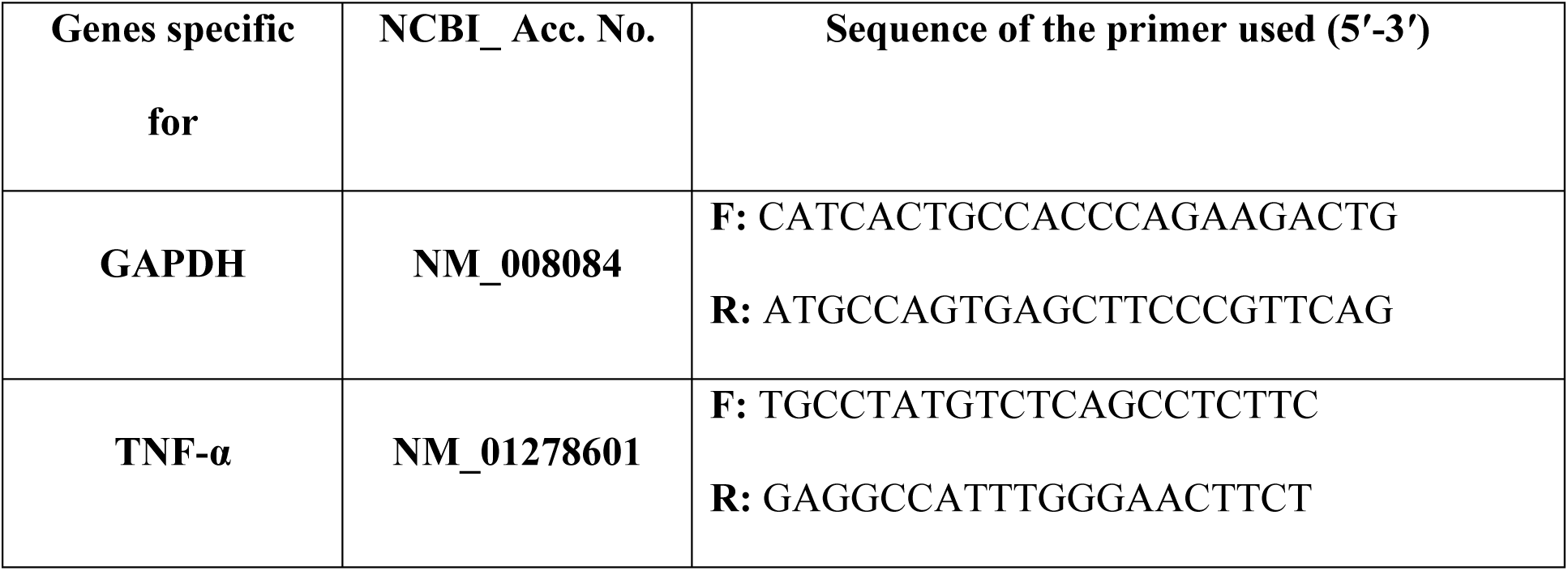
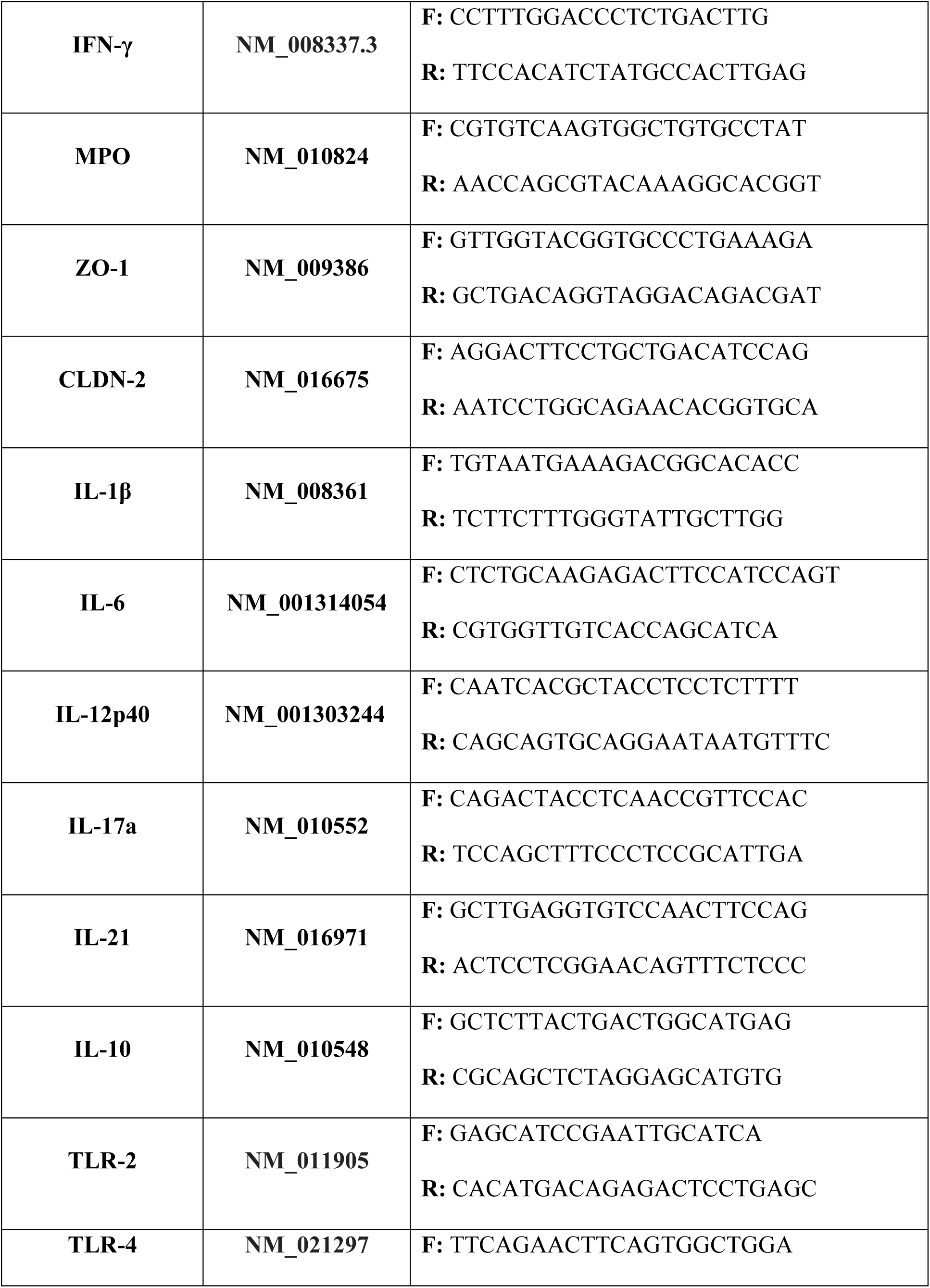
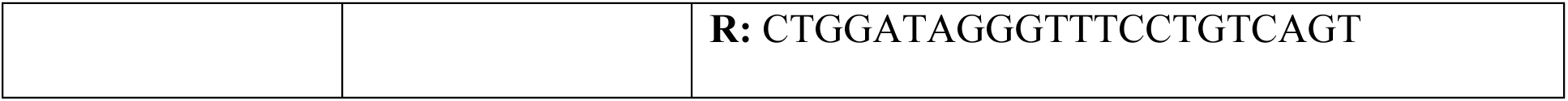
Sequences of forward (F) and reverse (R) primers used in gene expression studies.

### Myeloperoxidase (MPO) Activity Assay-

Colon tissues were collected from control and treated group of mice and rapidly homogenized in 4 volume of MPO assay buffer (5 g HTAB in 1 L of potassium phosphate buffer). To remove the insoluble debris tissue homogenates were centrifuged at 13000 rcf for 10 minutes at 4°C. After that the assay was performed as manufacturer protocol (Cat#MAK068, Sigma-Aldrich, St.Louis, USA) (18).

### C-Reactive Protein Assay-

Colon tissues were collected from control and treated group of mice and rapidly homogenized in tissue protein extraction buffer (Cat#78510, Thermo Scientific, Rockford, USA) containing 1X protease inhibitor cocktail (Cat#78429, Thermo Fisher Scientific, Rockford, USA). To remove the insoluble debris tissue homogenates were centrifuged at 13000 rcf for 10 minutes at 4°C. Protein concentration was normalized using Bradford assay (Cat#5000006, Bio-Rad, USA). The assay was performed according to the manufacturer protocol (Cat# ELM-CRP, Norcross, GA) (19).

### Lipocalin-2 (LCN2) Assay-

To perform this assay serum samples were collected from control and treated group of mice. Protein concentration was normalized using Bradford assay (Cat#5000006, Bio-Rad, USA). The assay was performed according to the manufacturer protocol (Cat# ELM-Lipocalin2, Norcross, GA) (20).

### Endotoxin Assay-

Endotoxin level was checked from the serum sample of both control and treated group of mice. Protein concentration was normalized using Bradford assay (Cat#5000006, Bio-Rad, USA). The assay was performed according to the manufacturer protocol (Cat#L00350, Piscataway, NJ, USA) (21).

### Detection of stool IgA-

IgA level was checked from cecal content of both control and treated group of mice. Cecal contents were dissolved in fecal immunoglobulin extraction buffer (1X PBS, 0.5%Tween, 0.05% Sodium Azide) containing 1X protease inhibitor cocktail (Cat#78429, Thermo Fisher Scientific, Rockford, USA) and centrifuged at 1500 rcf for 20 minutes at 4°C to isolate the supernatant. Protein concentration was normalized using Bradford assay (Cat#5000006, Bio-Rad, USA). The assay was performed according to the manufacturer protocol (Cat# ELM-IGA, Norcross, GA) (22).

## Data availability

All data generated or analysed during this study are included in this article.

## Acknowledgments

We are thankful to NISER animal house facility for assistance in preparing the slides for histo-pathological analysis.

## Author contributions

SM performed all experiments and drafted the manuscript. SM and PA designed the experiment. PA conceptualized, supervised the studies and finalized the manuscript.

